# Contribution of quasi-fibrillar superstructures in peroxide quenching by collagen peptides derived from fish processing by-products and their application as natural food additives

**DOI:** 10.1101/2021.02.11.430827

**Authors:** K Saleem, Pritha Dey, Charitha Sumeet, Mayur Bajaj, Y Geetika, A Vishwadeep, Pawan Tagadghar, Pradipta Banerjee

## Abstract

This study attempts to identify the significant role played by the secondary structure of collagen-derived peptides that are involved in lipid peroxide quenching in food products. Collagen was extracted from the skin of Perch and swim bladder of Rohu at 45-78% efficiency. It was identified as type-I based on a high molecular weight (110kDa) and its ion-exchange elution profile. The collagen samples were enzymatically hydrolyzed and collagen hydrolysate (CH) was extracted with an efficiency of 0.67-0.74g/g of collagen. The CH samples displayed a molecular weight in the range of 8.2-9.7kDa and exhibited a higher abundance of charges resulting in higher solubility. The structural studies revealed that the CH peptides existed in polyproline-II helix and formed a mimic-triple helix in a wide range of pH. In neutral and alkaline pH, the mimic helices joined to form a hierarchical quasi-fibrillar network that was smaller than collagen fibrils but also more dynamic. The CH exhibited >95% degradation in 15h through simulated digestion. The CH were able to decrease peroxide formation by 84.5-98.9% in commercially available cod liver and almond oil and increased the shelf life of soya bean oil by a factor of 5 after 6 months of storage. The addition of CH to cultured cells quenched peroxide ions generated *in situ* and decreased stressor activity by a factor of 12. The reason behind the high efficacy of CH was deciphered to be the proximal charge stabilization by the quasi-fibrillar network, which allowed efficient peroxide quenching and long-term stability.

## 1. Introduction

Lipid-based foods are prone to peroxidation, resulting in rancidity accompanied by decreased nutritional value and reduced shelf life [1]. The use of synthetic peroxide inhibitors as food additives effectively reduces peroxidation but is accompanied by health risk upon consumption [2-5]. This necessitates a search for natural peroxide quenchers that along with augmenting the shelf life of lipid-based food, also incorporates a substantial amount of nutritive content in the food products.

The marine industry is an active contender for the generation of numerous bioactive agents. The use of marine source-derived fatty acids, polysaccharides, proteins, and peptides has escalated in the last decade [6]. Collagen hydrolysates isolated from a large number of fishes including mackerel [7], seabass [8], salmon [9], yellowfin tuna [10], carp [11], croaker [12] and tilapia [13] display antioxidative activity. Even so, the major factors that contribute to this activity remain an actively researched topic with a substantial number of conflicting reports [6].

Broadly, the radical quenching properties of the peptides can be explained by two different doctrines. The first theory focuses on the importance of the amino acid residue sequences. In this school of thought, the secondary structure of a peptide does not play any major role, with the activity primarily depending on the presence of certain essential residues like Gly, Pro, Leu, Ala, Phe, Tyr, Trp, and His [14-16]. The alternate theory proclaims that the structural arrangement of the peptides plays a major role in imparting the peptide with bioactivity. However, major contradictions exist regarding the exact function of the secondary structure. Several studies have reported that peptides with a combination of low α-helix and high β sheet and/or β turn content display high bioactivity [17,18] and that the presence of random coil does not have a substantial contribution [18,19]. On the other hand, some studies have reported that peptide antioxidative property increases with the presence of random coil [20,21] and with the ability of peptides to self-assemble [21,22]. Very few studies have ventured into deciphering the exact role of secondary structure, if at all any, in enhancing antioxidative attributes. The contradictory results about the role of secondary structures and a lack of a mechanism for peroxide quenching create a lacuna. Bridging this gap is imperative as knowledge of the secondary structure’s role in radical quenching will allow the designing of novel peroxide quenching peptides with greater bioactivity.

The present study was conducted with a dual aim including utilization of fish skin and bladder-derived collagen hydrolysates as peroxide quenching agents in lipid-based food leading to shelf-life enhancement along with identifying the significance of peptide secondary structure in peroxide quenching.

Here, we report the first study on the characterization of quasi-fibrillar superstructures formed by peptides in collagen hydrolysate. We report that the fish by-product derived CH can have a peroxide quenching ability close to that of standard radical quenchers and can enhance the shelf life of foods over a prolonged duration. We show that this high efficacy can be traced back to the formation of a new kind of secondary structure termed as quasi-fibrils that is stable over a wide range of pH for effective radical quenching and dynamic enough to allow fragmentation by digestive enzymes.

## 2. Materials and Methods

### 2.1 Materials

Butyl-hydroxy toluene (BHT) was purchased from SD Fine Chemicals, Bangalore, India. Sephadex G100, Linolenic acid, and molecular weight markers, as well as rat tail tendon collagen, were brought from Sigma-Aldrich (St. Louis, MO, USA). Other chemicals used were of analytical grade. Cells were obtained from NCCS, Pune and media was procured from Himedia, Bangalore.

### 2.2 Collection of fish waste and extraction of collagen

Rohu and Perch derived waste were chosen to be recycled based on seasonal and market availability. The swim bladder of Rohu (RoSB) and skin of Perch (PeSk) were procured from the fish market near Kumarswamy Layout, Bangalore, Karnataka. Collagen type-I was isolated from the swim bladder and skin by acid extraction followed by salt precipitation [23]. The post-treatment residue (K4 g) was stirred in acetic acid for 48hrs at 4°C and filtered through cheesecloth to obtain soluble collagen. The insoluble fraction was dried and weighed (K5 g) and the weight of solubilized collagen (K6 g) was calculated. The % of soluble collagen was calculated as shown below.

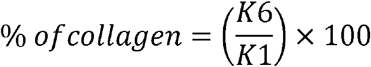

The collagen samples from Perch skin (PeSkC) and Rohu swim bladder (RoSBC) were precipitated with a 6% NaCl solution, filtered, and re-dissolved in acetic acid. The solutions were dialyzed in cellulose membranes with a molecular weight cut-off of 6kDa for 72hrs and lyophilized in a VD-250R Freeze dryer (Taitec, Japan).

### 2.3 Characterization of isolated collagen

#### 2.3.1 Electrophoretic pattern

The extracted collagen was subjected to SDS-PAGE with an 8% resolving gel following Laemmli’s [24] method and destained according to the method described by Wong et al. [25].

#### 2.3.2 FPLC Elution profile

PeSkC and RoSBC samples were applied to a 1.8×13cm Sephadex G100 column at a flow rate of 1ml min^-1^ using an ÄKTAprime plus unit (GE Healthcare). Void volume was calculated by running blue dextran. The column was calibrated by running standard markers ranging from 3kDa to 200kDa and the molecular weight of the collagen samples was calculated from the calibration curve.

For ion-exchange profiling, PeSkC and RoSBC were dissolved in 0.1M acetate buffer of pH 4.5 and applied to a 1.8×4.5cm CM Sepharose column at a flow rate of 1.2ml min^-1^. After 5min of buffer flow, a salt gradient of 0-1M NaCl was applied for a period of 30min.

### 2.4 Fragmentation of collagen

The dialyzed collagen samples were suspended in phosphate buffer and Clostridial collagenase type-I was added at a weight ratio of 300:1 (substrate: enzyme). The reactions were incubated at 37°C for 24hrs following its termination by freezing. The samples were lyophilized to obtain crude hydrolysate powder.

### 2.5 Purification of CH

The lyophilized materials were dissolved in 0.1M acetate buffer, pH 4.5, and injected in a 1.8×4.5cm column of CM Sepharose. The samples were eluted using 0.1M acetate buffer with a NaCl gradient of 0-1M for 25ml. Fractions representing peaks were pooled and applied to a 1.2×5cm desalting column. The desalted eluates were lyophilized and the dried powders were designated as RoSBCH and PeSkCH for Rohu swim bladder collagen hydrolysate and Perch skin collagen hydrolysate respectively. The yield was calculated as follows:

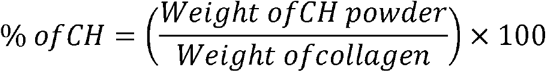

### 2.6 Characterization of CH

#### 2.6.1 Molecular weight profile of CH

The CH samples were dissolved in 0.1M acetate buffer, pH 4.5, and subsequently loaded into a 1.8×13cm Sephadex G100 column at a flow rate of 1mlmin^-1^. Elution buffer was run for 30min and 3ml fractions were collected. The column was calibrated as mentioned previously. The elution volume of the CH sample was noted and the molecular weight of the CH was calculated from the standard curve.

#### 2.6.2 Solubility profile of CH

The solubility profile of the CH was carried out as described by Tsumura et al. [26] by adding CH to buffers with pH ranging from pH 4 to 9 followed by centrifugation. The percentage of soluble fractions was calculated at each pH with the following equation:

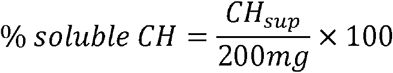

where *CH*_*sup*_ is the amount of CH present in the supernatant in mg. Solubility analysis was carried out in triplicate. Intact collagen was used as control.

#### 2.6.3 Structural analysis of CH

A Nanotrac wave II Q was used to measure the zeta potential and size of the CH samples dissolved in buffers with pH varying from 4-9. The Circular Dichroism (CD) spectra were obtained in two different buffer systems (pH 4 and 7) using a J-715 spectropolarimeter (Jasco, Japan). The functional groups of the CH samples were detected using Thermo Nicolet iS5 Fourier Transform-IR. XRD profiles of the CH samples were obtained using a Rigaku D/Max X-ray diffractometer with Cu Kα radiation source using a scan rate of 10° min^-1^ from 5° to 65°. Intact collagen was used as a control for all the structural studies.

#### 2.6.4 Bioavailability of CH

The CH was subjected to simulated digestion via the method of Song et al. [27]. A two-stage *in vitro* digestion system was adopted where the samples were subjected to 3h of pepsin treatment followed by 27h of pancreatic enzyme cocktail. Values obtained at the 0^th^ hour were taken to be 0% hydrolysis. Aliquots were aspirated at scheduled time intervals, treated, and analyzed for the presence of amino terminals. Intact collagen and 0.05M glycine solution were used as negative and positive control respectively.

### 2.7 Antiperoxide activity of CH

#### 2.7.1 Antiperoxide assay with linoleic acid and oils

The CH samples were assayed for their antiperoxide activity by using the protocol of Phosrithong and Ungwitayatorn [28] modified by Citta et al. [29] in a linoleic acid (LA) model with 10nM cumene hydroperoxide. LA supplemented with BHT and intact collagen were taken as the positive and negative control respectively. The antiperoxide activity of the hydrolysates was also tested against commercially available mustard oil (MO) and cod liver oil (CLO) following a similar protocol. Oil without added hydrolysate was used as control. The percentage of LPO reduction was calculated according to the equation:

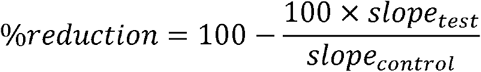

#### 2.7.2 Shelf-life enhancement assay

A shelf-life enhancement assay was performed to determine the effectiveness of the hydrolysates on a longer time scale. The assay was performed for a duration of 6 months on Soyabean oil (SBO) - a commercially available food product. For the test sample, RoSBCH was dissolved in 0.1M phosphate buffer, pH 7 at a concentration of 1.5mgml^-1,^ and mixed with 25% SBO in water: ethanol mixture. The control sample received only phosphate buffer. BHT mixed with SBO at a concentration of 0.1mgml^-1^ was taken as positive control and RoSBC at a concentration of 1.5mgml^-1^ was used as a negative control. Aliquots were aspirated once every week from each sample to test for LPO generation following the protocol described previously. The % reduction in LPO was calculated as before.

#### 2.7.3 Antiperoxide activity of CH in cell culture

Vero cell lines were maintained according to standard protocol [30] and the viability of cells was checked by trypan blue assay. The CH was dissolved in DMEM medium at a concentration of 1mgml^-1^ and added to 3.2×10^3^ cells in a sterilized 48 well microplate. A similar number of cells were cultured without the addition of hydrolysate. Cell adhesion and proliferation were checked at regular intervals through MTT assay with the help of a Bio-Rad 680 microplate reader. Absorbance obtained was correlated with a standard curve formulated between the number of live cells and absorbance of formazan dye.

CH-induced inhibition of LPO generation was checked according to the protocol given by Das et al. [23]. Briefly, cells were grown in 48 well plates to 80% confluency. The CH and fatty acids were added to the cells at a final concentration of 100mM (0.89mgml^-1^) and 120µM respectively. Cumene hydroperoxide was added to the dishes at final concentrations ranging from 0, 0.1, 1, and 10µM. The total volume of the medium was kept constant at 1.5ml. The control group contained cells seeded on dishes without any addition of fatty acids or cumene hydroperoxide. After 6h incubation, adherent cells were counted and the stressor concentration ensuing 50% cell survival was calculated.

### 2.8 Identification of peptides through mass spectroscopy

The CH samples were analyzed through LC-MS/MS for identifying abundant sequences. The CH was suspended in deionized water with 0.1% formic acid and 5% acetonitrile. The solution was injected into a Dionex Ultimate 3000 Micro-LC equipped with a Thermo Syncronis C18 column. The CH was eluted with 0.1% formic acid in LC-MS grade water. The eluants were transferred to a Bruker ESI-QUAD-TOF Mass spectrophotometer. Spectra were acquired at 30Hz with maximum mass cut off at 10kDa. A high S/N ratio was maintained to detect peaks with high intensity. The MGF file obtained was uploaded to Mascot Search Engine V 2.7.0.5 (Matrix Science, UK). The data obtained were searched in two databases: Swiss Prot and Contaminants.

### 2.9 Statistical analysis

All the assays were done in triplicates. Data was analyzed in Origin Pro 9. Probability values less than 0.05 were considered significant.

## 3. Result and Discussion

### 3.1 Extraction and purification of collagen

The total amount of soluble collagen obtained from PeSk and RoSB was found to be 45.99% and 78.86% respectively, as seen in Table 1. Previous authors have reported a wide range of collagen yields varying from 5.4% from the skin of common carp [31] to 51.9% from golden carp [32]. Swim bladders tend to have higher yields when compared to skin and bones, as their structure is primarily made up of pure collagen [33]. The extraction efficiency of rohu swim bladder collagen in the present study was, on average, higher than previous reports, which ranged from 46.5% [34] to 54.5% [35]. The overall high efficiency obtained in this study was due to the use of mild reagents coupled with repetitive extractions, which although time-consuming, yields better results.

**Table 1.**
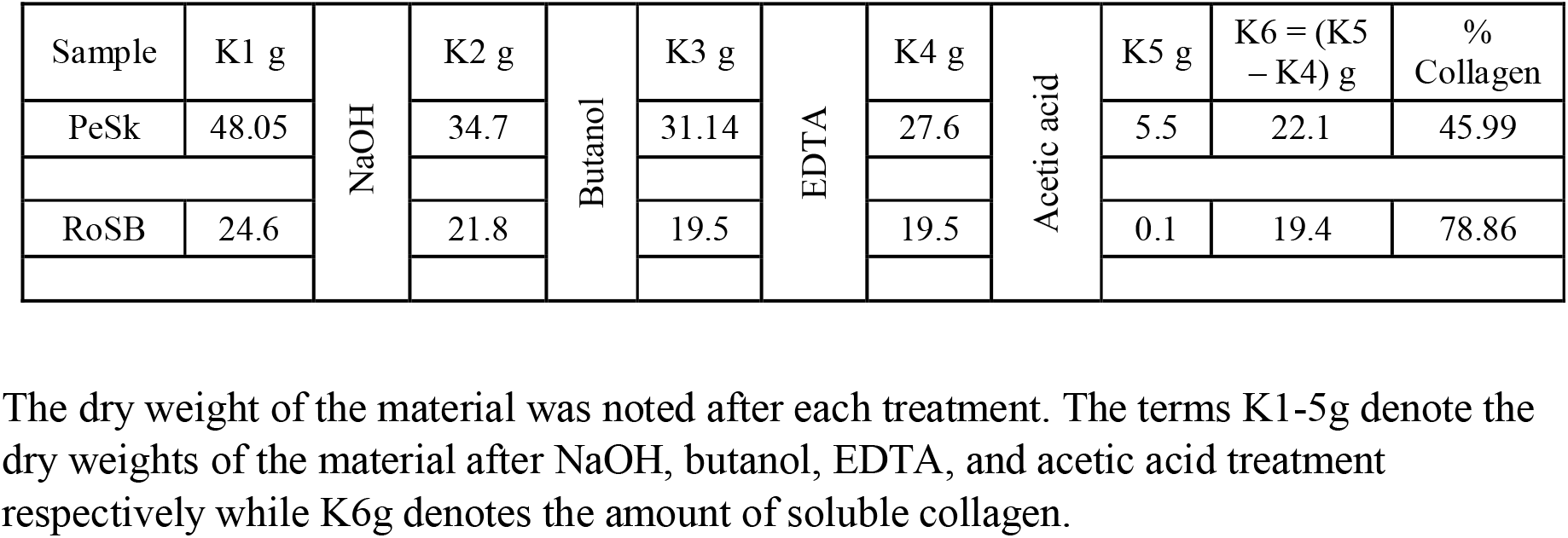
Extraction of collagen from fish waste.

| Sample | K1 g | NaOH | K2 g | Butanol | K3 g | EDTA | K4 g | Acetic acid | K5 g | K6 = (K5 – K4) g | % Collagen |
| --- | --- | --- | --- | --- | --- | --- | --- | --- | --- | --- | --- |
| PeSk | 48.05 |  | 34.7 |  | 31.14 |  | 27.6 |  | 5.5 | 22.1 | 45.99 |
| RoSB | 24.6 |  | 21.8 |  | 19.5 |  | 19.5 |  | 0.1 | 19.4 | 78.86 |

The dry weight of the material was noted after each treatment. The terms K1-5g denote the dry weights of the material after NaOH, butanol, EDTA, and acetic acid treatment respectively while K6g denotes the amount of soluble collagen.

### 3.2 Characterization of RoSBC and PeSkC

RoSBC and PeSkC displayed the specific band pattern of collagen with two α bands (α1 and α2) and high molecular weight β and γ bands as seen in Fig. 1a. The molecular weight of α1 and α2 chains were determined to be 118±2 and 100±2kDa respectively for both collagen samples. RoSBC and PeSkC eluted at 20.5min and 22.8min respectively in a Sephadex G100 column as seen in Fig. 1b. The single peak was due to the combined elution of α1 and α2 polypeptide chains of collagen type I. Substituting the V_t_/V_o_ value in the calibration graph, the molecular weight corresponding to the average α-chain was found to be 111kDa for RoSBC and 108kDa for PeSkC. The values correlated with the average molecular weights obtained from SDS-PAGE. Collagen type-I exists in a net positive charged state at acidic pH, leading to a preferential binding with the negatively charged CM sepharose column. RoSBC (Fig. 1c) and PeSkC (Fig. 1d) eluted at 16.1 and 19.7% of NaCl respectively from a CM sepharose column. The lower salt% was indicative of the lower abundance of positively charged amino acid residues in the concerned samples.

**Fig. 1.**
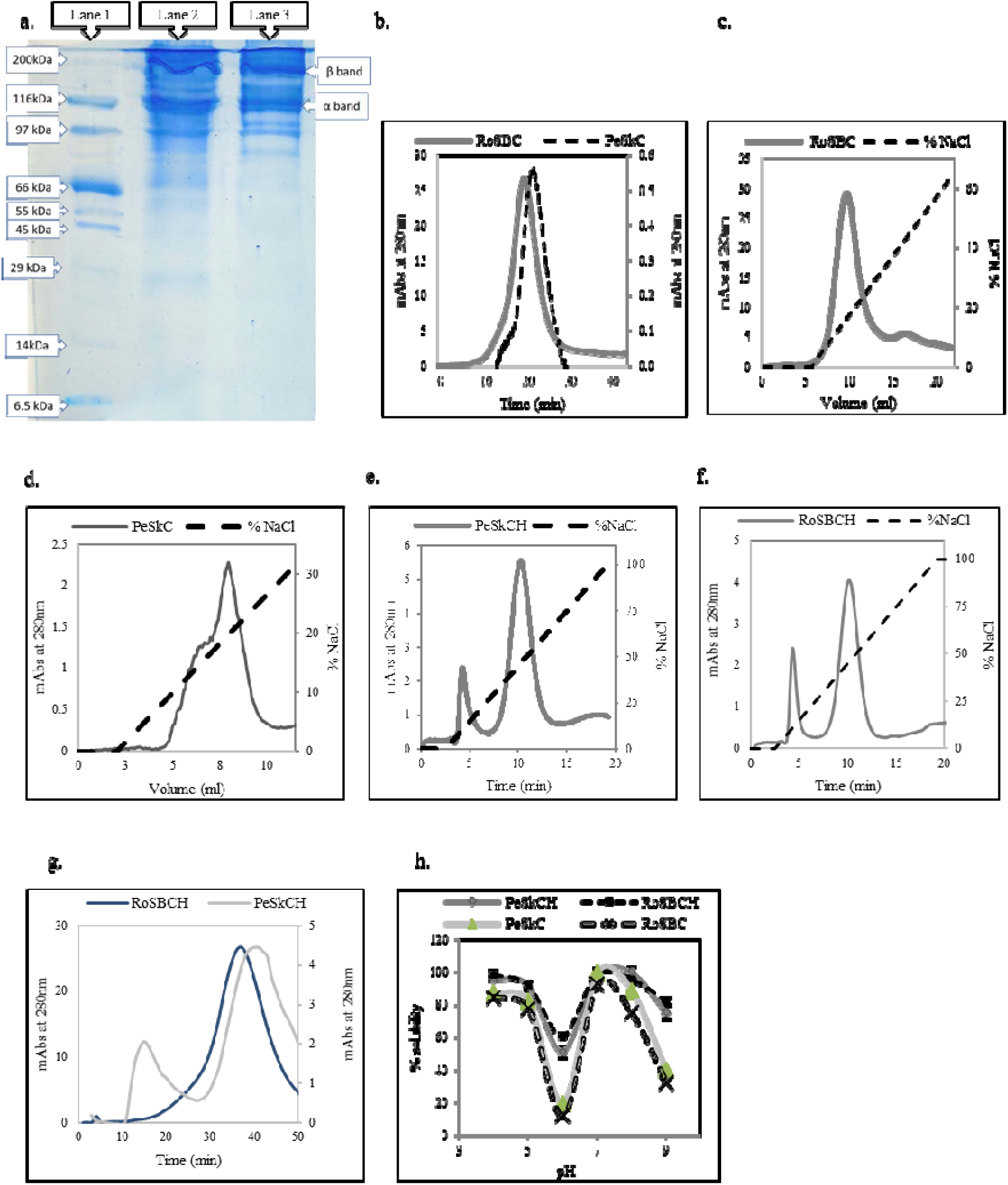
Characterization of collagen and collagen hydrolysate purified from Perch skin and Rohu swim bladder. a. SDS-PAGE of collagen samples; Lane 1, 2, and 3 indicates marker, PeSkC, and RoSBC respectively. a. Gel permeation chromatography elution profile of RoSBC and PeSkC. b. Ion exchange chromatography elution profile of RoSBC. c. Ion exchange chromatography elution profile of PeSkC. e. Ion exchange chromatography elution profile of PeSkCH. f. Ion exchange chromatography elution profile of RoSBCH. g.Gel permeation chromatography elution profile of RoSBCH and PeSkCH. h. Solubility profile of RoSBCH and PeSkCH compared to RoSBC and PeSkC in a pH range of 4 to 9.

### 3.3 Purification of RoSBCH and PeSkCH

As evident from Fig. 1e and 1f, the ion exchange elution pattern of PeSkCH and RoSBCH depicts two peaks: a large peak at 45±2% NaCl representing the free peptides and a smaller peak at 10.5% NaCl representing the coiled or aggregated peptides. The higher % of NaCl required to elute the CH stems from its increased charge density due to the larger number of free N and C terminal residues. Fig. 1g exhibits the elution profile of RoSBCH and PeSkCH. The major elution peak was at 37-41min in G100 and the molecular weight was determined graphically to be in the range 8.2-9.7kDa. The % yield of CH was calculated to be 0.67g/g of collagen and 0.74g/g of collagen for PeSkCH and RoSBCH respectively, which was better than previously reported results [8].

### 3.4 Solubility profile of RoSBCH and PeSkCH

The increased number of charges and smaller size improves the overall solubility of CH in various buffers. This increases its chances of being used as an additive in food items with a wide range of pH. The hydrolysate samples PeSkCH and RoSBCH displayed significantly similar (p>0.05; ANOVA) solubility patterns as depicted in Fig. 1h. However, collagen displayed a significantly different pattern (p<0.05, ANOVA) with low solubility at pH 6 and 9 due to their proximity to the isoelectric point of α1 and α2 chains respectively.

### 3.5 Structural analysis of RoSBCH and PeSkCH

#### 3.5.1 pH based DLS/ ZP profile

The DLS data given in Fig. 2a corroborated the solubility pattern. Both PeSkCH and RoSBCH exhibited similar structural hierarchy and surface charge. At pH 4 and 5, CH displayed a hydrodynamic radius of 145-173nm, which was smaller than collagen (270-292nm). The CH particle diameter was expected to be even smaller based on its low molecular weight. However, the peptides in CH were possibly coiling with each other to result in a larger size, a phenomenon distinctly noticeable during the ion exchange run. The α1 chain has its pI at 6 and therefore gains a neutral charge at the same pH, consequently, phasing out of the solution. The remaining chains, in turn, form smaller and possibly, incomplete helices. This decreases the average size of CH and collagen to 125±34 nm and 248±25 nm respectively.

**Fig. 2.**
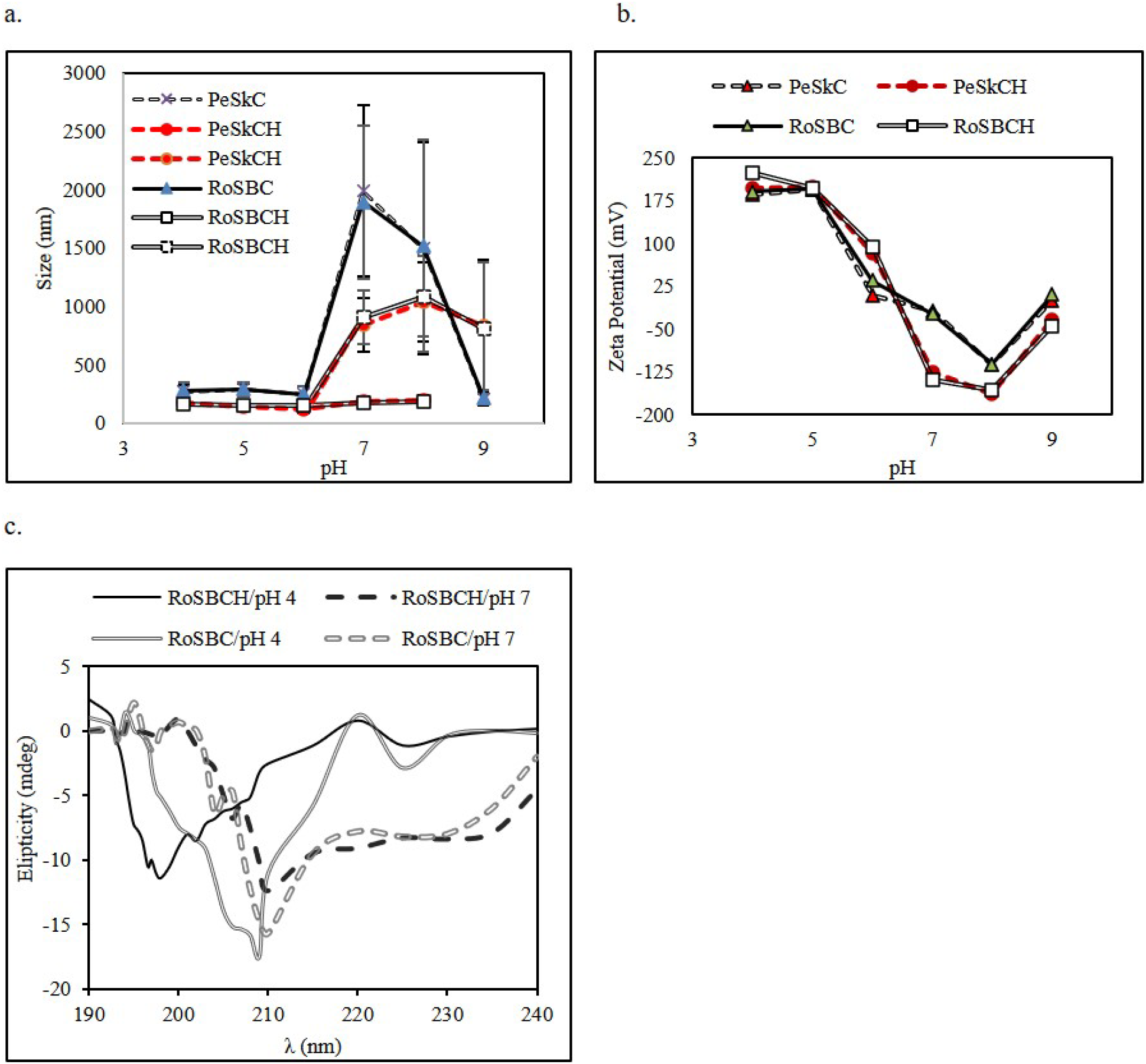
Structural characterization of CH. a. Size distribution of PeSkCH and RoSBCH compared to RoSBC and PeSkC in the pH range of 4 to 9. The CH samples split into two distributions in the neutral and alkaline range. Both are depicted in the graph with the same series title b. Zeta potential of PeSkCH and RoSBCH compared to RoSBC and PeSkC in the pH range of 4 to 9. c. CD spectral profile of RoSBCH at pH 4 and 7 compared to that of RoSBC under similar conditions. A similar profile was displayed by PeSkCH.

At alkaline pH, CH existed in two size distributions, one at 190±42nm and another at 842±229nm. The twin distributions indicated two levels of association in CH; one smaller, with regular coiled peptides and a second larger hierarchical form, that possibly resembles a fibrillar assembly. A similar situation was observed at pH 8. In the same pH range, collagen formed a large distribution at 1990±727nm indicating fibril formation. At pH 9, CH fully converted to this quasi-fibrillar form with an average size of 804±560nnm while, collagen lost the fibrillar hierarchy and reverted to smaller helices. This was because the α2 chain of collagen has a pI around 9 forcing it to phase out of the solution. The result was a deformed triple helix and consequent inhibition of the lateral associations required to form a fibril. Overall, this quasi-fibrillar assembly of CH was found to be more stable over a wide range of pH in comparison to that of collagen.

The zeta potential gives an idea of the stability of a biomolecule. A greater ZP, either positive or negative, indicates that the particles would repel each other and will not aggregate or precipitate out of the system [36]. As seen in Fig. 2b, the ZP profile of CH is significantly similar (p>0.05) to collagen in the acidic range. However, from pH 6 onwards, the ranges differ significantly (p<0.01). In pH 6, the ZP of CH is maintained at 83-94mV while collagen exhibits a ZP near 0, possibly owing to the denaturation of the α1 chain. In the neutral and alkaline range, CH maintains a ZP that was, in general, more negative than the ZP of collagen. A larger negative value was indicative of greater stability of the CH peptides in solution, in comparison to collagen. The ZP profile of CH matched with its size distribution and could be attributed to the dynamic but more stable quasi-fibrils of CH.

#### 3.5.2 CD profile of CH

Circular dichroism studies help in understanding the secondary structure of biomolecules based on their selective absorption of right or left circularly polarized light. The profile of RoSBCH and the control sample RoSBC is depicted in Fig. 2c. At pH 4, the CH and collagen exhibited a negative peak at 197nm and 209nm respectively, indicating the predominance of PP-II helices, which was expected given the greater abundance of imino acids in collagen. However, both CH and collagen also exhibited positive values at 222nm, indicating the formation of an ordered structure, possibly a triple helix [37,38].

At pH 7, the CD profile of the CH changed drastically. It showed a large minimum at 210nm and a negative plateau at 222nm, indicating that the ordered assembly had morphed into larger quasi-fibrillar helical structures. The control collagen sample displayed a similar profile, including a minimum at 210 nm and a negative plateau ranging from 220-230 nm, indicating fibrillogenesis [38].

It was interesting to note that the CD profile of CH mimicked that of collagen. This shift in the hierarchical state provided clinching evidence that the CH peptides did not exist in a random coil. At pH 4, the CH displayed smaller protofibrillar assemblies, while at neutral pH, it assembled into quasi-fibrillar structures, corroborating the solubility and the DLS/ZP profile.

#### 3.5.3 IR spectral profile of CH

The FTIR spectra of a protein is a powerful tool to investigate its secondary structure. The structural assembly of a polypeptide chain can be deduced by correlating various vibrating frequencies of its peptide bonds to specific hydrogen-bonding patterns that characterize myriad secondary structural elements [39].

Fig. 3a and 3b display the IR profile of PeSkCH and RoSBCH respectively. The Amide A band (3200-3400cm^-1^) is generally associated with the peptide bond N-H and its propensity to participate in hydrogen-bonding. The presence of the band at 3282-99cm^-1^, the low wavenumber end of Amide A sub spectrum, indicated the presence of intra-or inter-chain hydrogen bonding [40]. PP-II chains in triple helices are interconnected by one hydrogen bond per helical rise. This low frequency of hydrogen bonding is reflected in the slightly lower intensity of Amide A. The Amide B band, arising due to the CH_2_ asymmetric stretching, was visible at 2924-36cm^-1^.

**Fig. 3.**
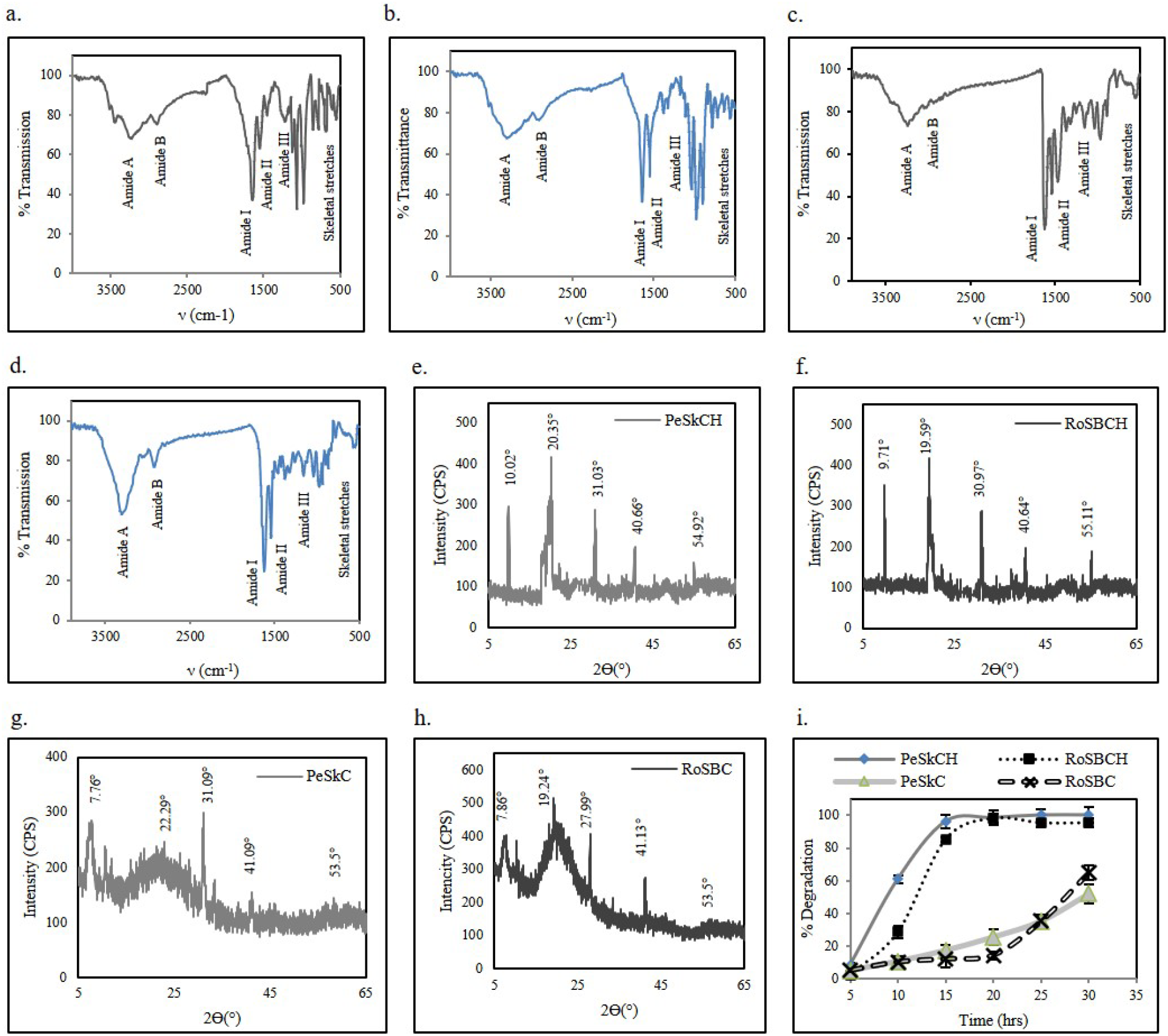
FTIR, XRD and digestion profile of CH. FTIR profile for a. PeSkCH b. RoSBCH c. PeSkC d. RoSBC; XRD profile for e. PeSkCH f. RoSBCH g. PeSkC h. RoSBC i. In vitro digestion study on PeSkCH and RoSBCH with PeSkC and RoSBC as controls. The digestion profiles for collagen and CH samples were significantly different (p<0.01).

The Amide I band (1610-1680cm^-1^) is affected in specific ways depending on the environment of the peptide bond and it provides sufficient clues that lead to the identification of the secondary structure [41]. A prominent peak in the sub-range of 1635-41cm^-1^ represented the presence of PP-II helices [42]. The Amide II band (1520-1580cm^-1^) occurs due to the C-N stretching in combination with N-H bending and its variations can reveal the tertiary structure of a biomolecule. Both CH samples displayed a peak at the range of 1540-45cm^-1^ confirming the presence of triple helices [43]. However, the bands were of lower intensity when compared to the control collagen samples.

A prominent peak at 1451-6cm^-1^ and a smaller band at 1336-8cm^-1^ were attributed to the wagging of proline CH_2_ and the presence of side chain vibrations, respectively [44]. The Amide III bands displayed a peak at 1202cm^-1^ and a shoulder peak around 1244cm^-1^. The former was attributed to the vibration of amino acid side chains in a triple helix and the latter was attributed to inter-strand hydrogen bonding in a distorted triple helix [39]. The high-intensity peaks at 960, 1058, and 1123cm^-1^ occurred due to the skeletal stretching of the peptides. The presence of GPO triplets confers endo-exo imino ring pucker motions that, coupled with the peptide backbone, leads to expansion and compression of the regions surrounding the triplets [45].

Fig. 3c and 3d depict the IR profile of control samples PeSkC and RoSBC respectively. The samples displayed a prominent Amide A at the range of 3285-7cm^-1^, indicating the stability of helices via hydrogen bond formation. Strong amide I, II, and III bands were visible at 1637-41, 1547-50, and 1238-9cm^-1^ respectively were attributed to the presence of abundant PP-II and triple helix. Prominent bands at 1450-1 and 1336-8cm^-1^ confirmed the presence of proline CH_2_ and side-chain vibrations in collagen respectively [44]. The stretches around 1400-1450 and 983-1058cm^-1^ corresponded to carbonates and skeletal stretches arising due to GPO triplets respectively.

To summarize, the CH peptides displayed a propensity to form PP-II helices that assembled in a distorted triple-helical manner. However, the assembly was less robust than collagen due to the smaller size of the peptides, as evident from the prominent GPO skeletal stretches in CH.

#### 3.5.4 XRD profile of CH

The diffraction of incident X-rays by biomolecules can lead to an understanding of their structural organization. Fig. 3e and 3f portray the XRD profile of PeSkCH and RoSBCH respectively. The band at 9.71-10.02° denoted an inter-planar distance of 0.883-0.911nm, which corresponded to the equatorial distance between the PP-II chains in a triple helix [46]. The smaller intermolecular packing distance, indicated that the helix formed by the peptides were of smaller proportions compared to the triple helix of collagen. The peak at 19.59-20.35° corresponded to an interplanar distance of 0.425-0.453nm. The distance was slightly higher than the regular amorphous scattering distance of tropocollagen units and corroborated the existence of deformed triple helices in CH. A third peak was observed in 30.97-31.03° with an interplanar distance of 0.295-0.319nm. This matched with the helical rise per residue in single peptide chains when oriented in a triple helix [47]. The peak at 40.64-40.66° represented a distance of 0.219-0.22nm, which matched with the helical rise per residue near the peptide chain extremities, particularly, non-helical regions [48]. The peaks at 54.92-55.11° represented a length of 0.171nm, which was close to the radius of a single PP-II helix [49].

The control collagen samples displayed typical peaks as displayed in Fig. 3g and 3h. The first peak was observed at 7.76-7.89°, with an interplanar distance of 1.125-1.139nm, the usual distance between PP-II chains in a triple helix [46]. The second peak was diffused in nature for both collagen samples, ranging from 18-24° arising due to the diffraction of X-rays from multiple tropocollagen-like units, scattered in myriad directions [46]. The peak at 53.5° confirmed the presence of the PP-II helix and the peaks near 27.99-31.09° and 41.09-41.13° indicated the presence of residues in the helical and terminal regions of the PP-II helix respectively.

To summarize, the peptides in CH had a distinct PP-II conformation with helical mid-regions and deformed-helical extremities. Each PP-II chain coiled with other chains to form a “mimic” triple helix, that was smaller in dimensions than the triple helix of collagen.

### 3.6 Digestion profile of RoSBCH and PeSkCH

The simulated digestion assay provides the rate of clearance of the ingested compounds by the digestive system. Ideally, a bioactive agent added to a food should be removable by digestive proteases before it reaches circulation and liver, consequently reducing the chances of bioaccumulation. As seen in Fig. 3i, both the hydrolysates exhibited significantly similar (p>0.05, ANOVA) fragmentation profiles and were markedly different from that of the intact collagen samples (p<0.05, ANOVA). The CH displayed >95% degradation after 15hrs while the collagen samples were only 50-60% degraded by 24hours. As mentioned before, the triple helically enforced fibrillar structure of collagen imparts low protease susceptibility resulting in decreased digestibility coefficient [50,51]. CH samples had a quasi-fibrillar structure which was partially similar to that of collagen but less sterically congested. This rendered it vulnerable to enzymatic digestion resulting in faster fragmentation. The free amino acids obtained as the products of digestion could be incorporated by the consumer in their amino acid pool leading to additional benefit. Overall, the results confirm that upon ingestion, a major portion of CH was removable by digestion.

#### 3.7 Lipid Peroxidation Inhibition Assay

Lipid-rich food products generate peroxyl ions upon being stored for a longer duration, even with necessary packaging precautions, consequently leading to various health problems upon consumption. The LPOI assay simulates the longer storage period by the addition of catalytic amounts of cumene hydroperoxide (hereafter referred to as cumene), a lipid radical generating compound that hastens the peroxyl formation process [52]. The resultant peroxide oxidizes Fe^2+^ to Fe^3+^ which reacts with thiocyanate to produce colour. As shown in Fig. 4a, the control sample containing only linoleic acid displayed increasing peroxide levels with time. However, linolenic acid mixed with the hydrolysates showed a significant (p<0.05) reduction in lipid peroxidation. PeSkCH and RoSBCH reduced peroxide formation by 97-98%, a value significantly similar to that of BHT (p>0.05).

**Fig. 4.**
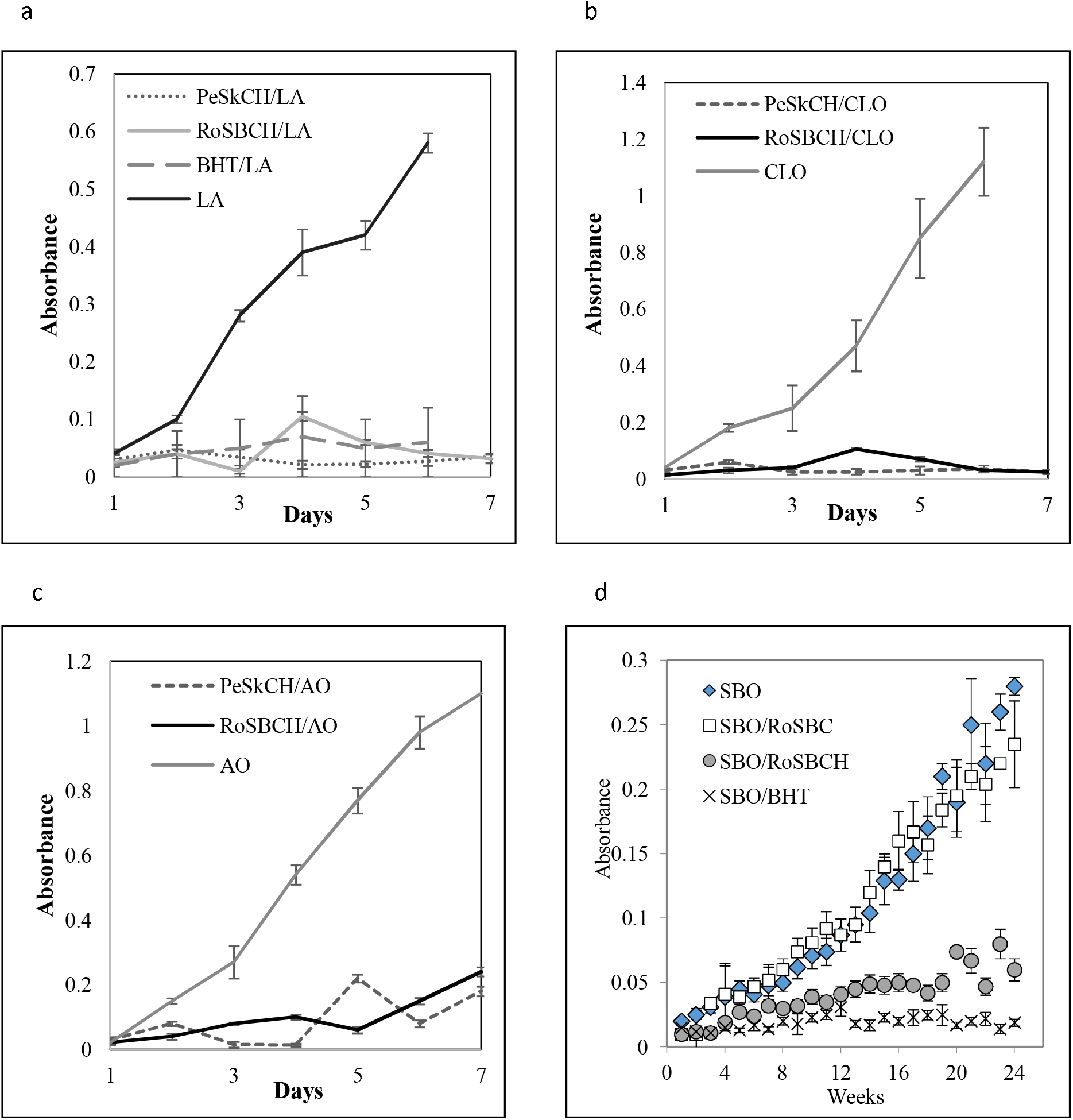
In vitro peroxide quenching activity of RoSBCH and PeSkCH. The hydrolysate sample displayed significantly different values (p<0.01) from the control sample. a. Antiperoxide activity of PeSkCH and RoSBCH with linolenic acid as control. b. Antiperoxide activity of PeSkCH and RoSBCH in cod liver oil. c. Antiperoxide activity of PeSkCH and RoSBCH in almond oil. d. Shelf-life enhancement of soya bean oil with RoSBCH for 6 months.

The assay was repeated with commercially available products - Almond and Cod liver oil as test samples. Both contain a mixture of mono and polyunsaturated fats like oleic acid, linoleic acid, ricinoleic acid, and palmitic acid [53,54] and are susceptible to peroxide formation upon storage. It was observed from Fig. 4b and 4c, that the control oil samples (without CH) displayed an increasing peroxide accumulation over the duration of the study. However, the samples with PeSkCH and RoSBCH could reduce cod liver oil peroxidation by 98-99% and almond oil peroxidation by 84-85% over a period of six days.

The significant resemblance in activities (p>0.05) of the hydrolysates was due to the presence of the same collagen type-I in all the samples, which upon fragmentation, generated peptides with closely resembling sequences. However, the minor variations within the values may be attributed to the existence of minor changes in amino acid sequences. The fact that lower molecular weight peptides or hydrolysates of proteins can produce better anti-oxidative properties has been established before [55]. Apart from CH, hydrolysates from hen egg lysozyme and quinoa have been also shown to exhibit lipid peroxide inhibition by 36.8-61% and 75.15% respectively [56,57]. Hydrolysates obtained by gastric and gastrointestinal digestion of proteins obtained from *Amaranthus caudatus* were able to reduce the lipid peroxidation by 72.86% and 95.72% respectively [58].

### 3.8 Shelf Life Enhancement Assay

Shelf life of a food is defined as the time duration for which, a product may be stored without becoming unsuitable for consumption. Environmental factors such as exposure of food components to heat, light, moisture, mechanical stress, and contamination can accelerate the degradation of food components leading to a decrease in shelf life [59]. Soyabean oil contains over 80% unsaturated fat, with a prevalence of linolenic acid, which increases the chances of peroxide formation upon storage [60]. As seen in Fig. 4d, the control samples with only SBO exhibited an increasing LPO profile over the period of study. The control SBO/RoSBC showed a similar profile (p> 0.05) indicating the inefficiency of intact collagen to quench peroxides. The PC, SBO/BHT, and the test sample, SBO/RoSBCH, were able to lower the LPO levels by 86% and 81.3% respectively. Overall, the CH was able to increase the shelf life of SBO by a factor of 5. The long-term efficacy displayed by the RoSBCH can be attributed to the stability of the quasi-fibrillar assembly.

### 3.9 Lipid Peroxidation Inhibition Assay in cell culture

Since the CH used in this study has to be incorporated into a food product, it has to be ensured that the additive does not have any ill effects on cells. As shown in Fig. 5a, Vero cells grown in the presence of RoSBCH and PeSkCH could adhere and proliferate without any significant difference (p>0.05, by ANOVA) when compared to that of control cells (without any CH).

**Fig. 5.**
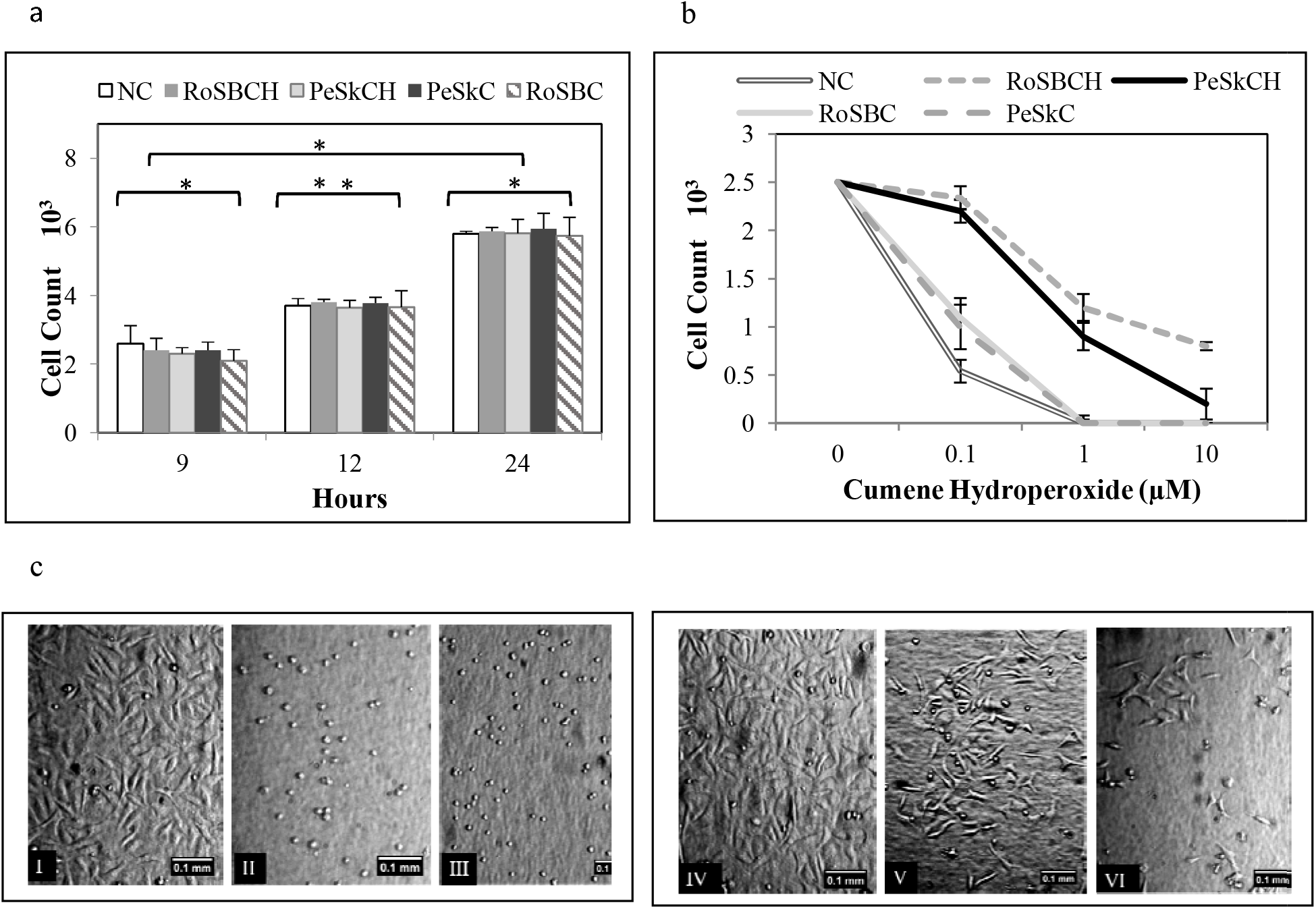
Cytoprotective and antiperoxide activity of PeSkCH and RoSBCH in mammalian cell culture. a. Vero cell adhesion/proliferation profile in the presence of PeSkCH and RoSBCH in culture medium, * - p>0.05 and ** - p<0.05. b. Antiperoxide activity of PeSkCH and RoSBCH in a cumene hydroperoxide/unsaturated fatty acid medium. c. Photomicrographs depicting the effect of LPO on cell adhesion and viability. The white line at the bottom right corner represents a line of 0.1mm. I, II, and III indicate control cells at 0, 1, and 10µM of cumene hydroperoxide respectively while IV, V, and VI indicate test cells at 0, 1, and 10µM of cumene hydroperoxide in the presence of RoSBCH.

As seen in Fig. 5b, the control cells were unable to sustain themselves at 0.1µM cumene and were seen to perish at a concentration of 1µM cumene. However, the presence of the CH had a significant effect (p<0.05) on cell survival rate. At a concentration of 1µM cumene, RoSBCH and PeSkCH could retain 48% and 36% of cells. RoSBCH was able to retain 32% viable cells even at a concentration of 10µM cumene. In the presence of CH, the LD_50_ of cumene hydroperoxide was increased by a factor of 12, validating its role as a cytoprotective agent. The intact collagen samples used as negative controls were unable to display any cytoprotection. Fig. 5c displays the photomicrographs depicting the effect of LPO on the cells. The control cells (I-III) exhibit gradual cell detachment and dead cells. However, the test cells (IV-VI) evidently show the presence of attached, live cells. Interestingly, the adhesion pattern of the cells changed during stress, as they became disarrayed and rearranged themselves.

To conclude, the hydrolysates were proven to be non-cytotoxic and were able to quench free radical generation in mammalian cell culture.

### 3.10 Identification of abundant peptides in PeSkCH and RoSBCH

Fig. 6a and 6b depict the LC-tandem mass spectroscopy profiles of PeSkCH and RoSBCH respectively. Collagen type-I is evolutionarily conserved, mostly comprising of repetitive sequences at high abundance. Consequently, the physicochemical behavior of the CH depends primarily on the structural attributes and the amino acid composition of these abundant sequences. Each peak observed in the figures is a conglomeration of masses of closely matching sequences with the most abundant peptides giving rise to the most intense peaks. Table 2 lists 16 sequences found in increased abundance in the two CH samples. The size of the peptides ranged from tetrapeptide to larger peptides with 33 and 47 residues. The peptide sequences followed the uniquely distinctive collagen signature of the Gly-Pro-Y repeating unit, where Y is often a Hyp residue. Gly was the most prevalent residue among the peptides with an abundance of 36% followed by Pro at 31.2%. The third most common residue was Ala at 9.28% followed by Arg (6.6%), Asp (5.1%), Gln and Val (2%), Glu, Lys, and Phe (1.5%) along with Hyp and Asn (0.5-1%).

**Table 2.**
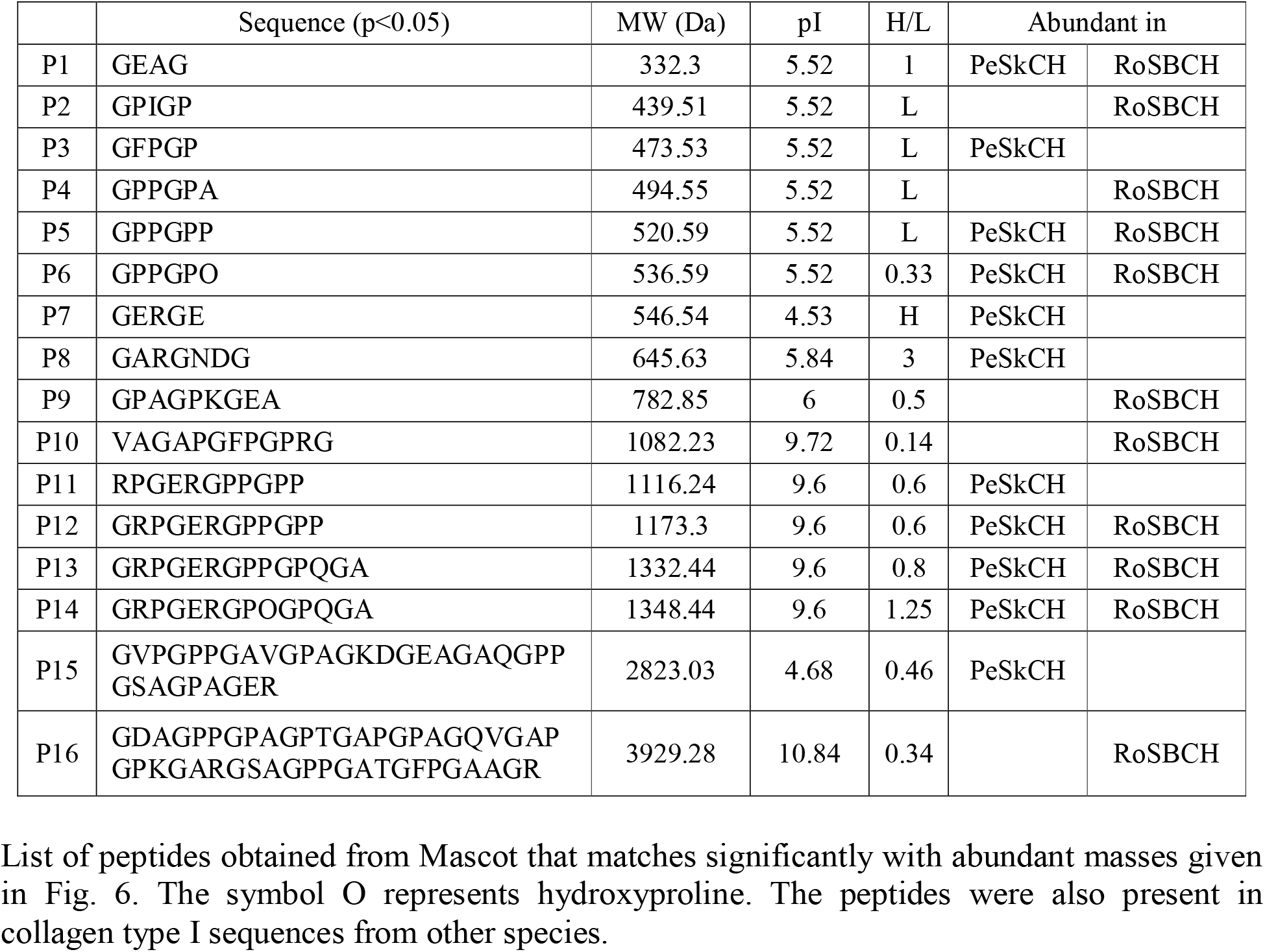

|  | Sequence (p<0.05) | MW (Da) | pI | H/L | Abundant in |  |
| --- | --- | --- | --- | --- | --- | --- |
| P1 | GEAG | 332.3 | 5.52 | 1 | PeSkCH | RoSBCH |
| P2 | GPIGP | 439.51 | 5.52 | L |  | RoSBCH |
| P3 | GFPGP | 473.53 | 5.52 | L | PeSkCH |  |
| P4 | GPPGPA | 494.55 | 5.52 | L |  | RoSBCH |
| P5 | GPPGPP | 520.59 | 5.52 | L | PeSkCH | RoSBCH |
| P6 | GPPGPO | 536.59 | 5.52 | 0.33 | PeSkCH | RoSBCH |
| P7 | GERGE | 546.54 | 4.53 | H | PeSkCH |  |
| P8 | GARGNDG | 645.63 | 5.84 | 3 | PeSkCH |  |
| P9 | GPAGPKGEA | 782.85 | 6 | 0.5 |  | RoSBCH |
| P10 | VAGAPGFPGPRG | 1082.23 | 9.72 | 0.14 |  | RoSBCH |
| P11 | RPGERGPPGPP | 1116.24 | 9.6 | 0.6 | PeSkCH |  |
| P12 | GRPGERGPPGPP | 1173.3 | 9.6 | 0.6 | PeSkCH | RoSBCH |
| P13 | GRPGERGPPGPQGA | 1332.44 | 9.6 | 0.8 | PeSkCH | RoSBCH |
| P14 | GRPGERGPOGPQGA | 1348.44 | 9.6 | 1.25 | PeSkCH | RoSBCH |
| P15 | GVPGPPGAVGPAGKDGEAGAQGPP<br>GSAGPAGER | 2823.03 | 4.68 | 0.46 | PeSkCH |  |
| P16 | GDAGPPGPAGPTGAPGPAGQVGAP<br>GPKGARGSAGPPGATGFPGAAGR | 3929.28 | 10.84 | 0.34 |  | RoSBCH |

**Fig. 6.**
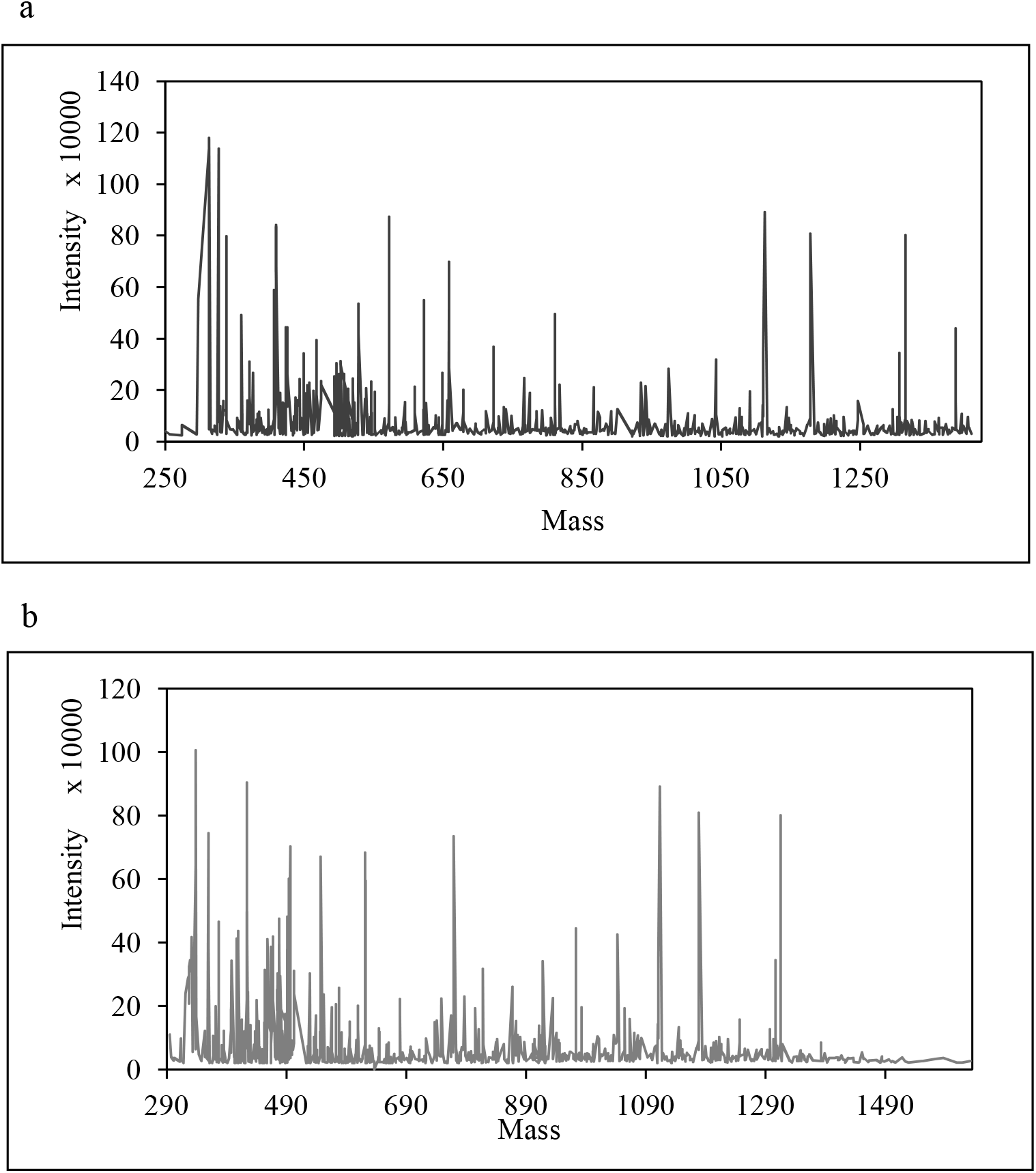
LC-Tandem Mass spectroscopy profile of a. PeSkCH b. RoSBCH displaying high-intensity peaks from abundant peptides.

The hydrophile to lipophile ratio (H/L) of a peptide is a good measure of the degree to which it can interact with a hydrophilic or a lipophilic medium. Based on the H/L ratio, the abundant peptides could be segregated into three classes. Class I comprised of the peptides P2-P6, P9, P10, P15, and P16, with a H/L ratio ranging from completely lipophilic to 0.5. Peptides in this group were responsible for the high lipid solubility displayed by CH. Class II comprised of peptides P11-13, displaying a H/L range of 0.6-0.8. Class III comprising of peptides P1, P7, P8, and P14 had the highest H/L ratio ranging from 1 to completely hydrophilic. These peptide classes were responsible for the high solubility of CH in a wide array of aqueous medium and their constituent amino acids were able to partake in radical quenching reactions.

### 3.11 Activity correlation with sequence and structure

General antioxidative peptides stabilize free radicals by the donation of a hydrogen atom or chelation of free metal ions. Gly was the most abundant residue and it has been reported to take part in radical quenching by donating the single hydrogen atom acting as its functional group [14,61]. Pro was present at higher abundance in the peptides when compared to collagen. The pyrrolidine ring of Pro residues can physically quench singlet oxygen species generated by lipid peroxides by forming a charge transfer complex [62,63]. Pro residues in collagen are generally unable to display this activity because of the rigid triple-helical structural constraint. The peptide classes with high H/L displayed a higher abundance of Asp and Glu residues, which are known to chelate catalytic metal ions or transfer H^+^ to radicals leading to quenching of lipid peroxides [64,65]. Hydrophobic amino acid residues like Ala and Val have also been reported to possess radical scavenging activity as the non-polar functional groups have been shown to react with hydrophobic chains of PUFA [66,67].

A major problem that most antioxidative agents suffer from is charge neutralization. After the donation of the hydrogen, the donor residues impart a negative charge which unless neutralized could possibly inhibit further H donation. Positively charged residues can offer the required stabilization by cation-pie interaction or through hydrogen bond [68]. However, this necessitates the positively charged group to be present within interacting distance and to be oriented in a manner suitable for the charge stabilization.

It is here that the structural characteristics of the hydrolysate play a major role. As seen in this study, the prevalence of Pro confers a PPII secondary structure to the peptides. The resulting sterical constraint forces the functional groups of the residues to be extended away from the backbone and towards the solvent, leading to their exposure to neighboring PPII chains [69,70]. As seen from the structural studies, three PPII chains wind around each other to create a mimic-helix with an average interchain distance of 0.9nm. The subsequent lower concentration of residues in the PPII conformational space lowers the entropy of the mimic-helix when compared to collagen and facilitates inter-helix associations leading to the formation of a large fiber-like hierarchical assembly [69]. The hierarchy of the resultant superstructure was retained over a wide range of pH due to the resilience of the PPII conformation. However, the presence of large skeletal stretches in FTIR combined with lower dimensions of the mimic helix indicated that the assembly was physically more dynamic than collagen and as such, was a “quasi-fibril”. This quasi-fibril works like a massive network allowing the amino acid residues of individual peptides to interact with the lipid peroxides and also to stay connected to each other through weak interactions. Such a structural constraint ensures that peptides with similar H/L values remain in close proximity. This confirms that the positively charged residues remain in a certain orientation that allows them to neutralize the charge developed on a neighboring H donor residue. Thus, any charge developed in one peptide is delocalized on to the entire fibrillar structure, essentially making the fibril a charge distribution network.

The results obtained in this study can help decipher previous apparently contradictory results. The fact that a structural hierarchy of CH quasi-fibrils orients functional groups and works as a charge delocalization tool making it more effective than any single antioxidant peptide, provides a suitable explanation to the results of two recent studies which state that antioxidative activity increases with peptide self-assembly [21,22]. It is quite likely that some of the peptides in other studies can form similar self-hierarchical structures while appearing as random coils in CD, explaining the relationship between high activity and prevalence of random coils, as reported by [20].

Collagen, on the other hand, displayed a strict fibril formation at pH 7. The dense coiled-up environment within the fibrillar structures creates sterical hindrances, preventing the functional groups from reacting with the peroxides [37,71]. Compared to this, the dynamic structure of the CH quasi-fibril allows the peptide functional groups the stability to retain their orientation while remaining flexible. Overall, the study also confirms the “structural” theory in which the secondary structure of the peptide plays an important role in the long term antiperoxide activity of the peptides.

## 4. Conclusion

Collagen type-I comprises 90% of the protein in most fish processing by-products and it can be extracted through several techniques. The high yield of collagen obtained in this study was due to the use of mild conditions and repeated extractions. The collagen was identified to be type-I based on its elution and electrophoretic profile followed by degradation with collagenase to obtain CH. Structural analysis of the CH revealed that the peptides existed in PPII conformation. In both acidic and alkaline ranges, the PPII peptides coiled into mimic helices, which, at neutral and alkaline conditions, further assembled to form large quasi-fibrillar superstructures. Compared to collagen fibrils, the quasi-fibril arrangement was less susceptible to pH changes but vulnerable to digestive proteases because of its higher dynamicity and smaller sizes of the constituent peptides. The CH was successfully able to reduce LPO formation in a standard model by 97-98% and up to 84.5-98.9% in almond and cod liver oil, along with enhancing the shelf life of soya bean oil after 6 months of storage. The CH samples were non-toxic and offered protection to mammalian cells faced with lipid peroxyl insult. 16 abundant peptides were obtained from the CH samples and they could be grouped into three categories based on their H/L ratio. The high abundance of Gly, Pro, Ala, Arg, and Asp indicated that LPO quenching occurred primarily through hydrogen transfer to quench the radical. The quasi-fibrillar network plays a significant role in allowing peptides to remain oriented at a reacting distance of each other so that the residual charge developed on the donor peptide can either be neutralized by a neighboring residue or delocalized in multiple peptides.

Based on the high efficiency of extraction, the *in vitro* radical scavenging activity, the higher stability of CH, and the known immune-compatibility of collagen peptides, it can be safely assumed that collagen hydrolysate from Perch skin and Rohu swim bladder could be a safe and improved alternative product to be added to lipid-based foods to increase their shelf life.

## Declarations

## Acknowledgments

The authors express their gratitude to the LCMS facility, Indian Institute of Science, Bangalore, India, for helping with the sequence identification. The authors also thank the Department of Chemistry at Dayananda Sagar University, for the help provided in using the FT-IR instrumentation facility.

## Funding

The authors are thankful to the Centre of Innovation in Science and Engineering in Dayananda Sagar Institution, funded by the Vision Group on Science and Technology, Government of Karnataka (Grant number GRD 316) for partial funding of this study. The authors also express their gratitude to the Department of Science and Technology, Govt. of India, (Grant ECR/2016/001526) for cell culture facilities.

## Competing interest

The authors declare that they do not have any conflicts of interest that are relevant to the content of this article.

## Ethics approval

Not applicable

## Consent to participate

Not applicable

## Consent for publication

Not applicable

## Availability of Data and materials

Not applicable. All the relevant data has been displayed in the manuscript figures.

## Authors’ contributions

Saleem K, Pritha Dey, Charitha Sumeet, and Mayur Bajaj were responsible for conducting decisive experiments, collection of data and its analysis, preparation of graphs, manuscript draft preparation, smooth running of the project including managing funds for the project. Geetika Y, Vishwadeep A, and Pawan Tagadghar worked on structure and sequence data of the peptides along with editing later drafts of the manuscript. Dr. Pradipta Banerjee was responsible for the conceptualization of the project, acquisition, and allocation of funds for the project, devising experimental schemes in discussion with the authors, statistical analysis, data interpretation, supervising the progress of the project along with manuscript writing and editing.

## Notes

### Competing Interest Statement

The authors have declared no competing interest.

